# Identification, quantification, and elimination of barcode crosstalk in multiplexed Oxford Nanopore sequencing

**DOI:** 10.1101/2025.11.19.689316

**Authors:** Qing Dai, Claudia K. Gunsch, Joshua A. Granek

**Affiliations:** Civil and Environmental Engineering, Pratt School of Engineering, Duke University, Durham NC, 27708, USA; Department of Biostatistics and Bioinformatics, Duke University School of Medicine, Durham NC, 27708, USA

## Abstract

Barcode crosstalk is a potential source of error in indexed multiplex sequencing that can mimic cross-contamination and generate false-positive signals, particularly in experiments with low-biomass samples and imbalanced designs. Here we identify and quantify barcode crosstalk in multiplexed Oxford Nanopore sequencing and introduce post-ligation pooling (PLP). PLP is a drop-in library-preparation modification that prevents barcode crosstalk, rather than simply mitigating its effects. These results highlight the potential for barcode crosstalk to have detrimental effects in multiplexed long-read sequencing and establish PLP as an immediately adoptable mitigation, especially where negative controls and low-abundance signals inform biological interpretation.

## Introduction

Index barcodes enable multiplexing in high-throughput sequencing experiments by allowing multiple samples to be pooled and sequenced within a single run. Multiplexing lowers per-sample costs and, when designed appropriately, helps manage batch effects by processing samples under shared experimental conditions. However, barcoding introduces the potential for read misassignment, in which reads are attributed to the wrong sample. Misassignment arises from two major sources: demultiplexing error and barcode crosstalk (also known as barcode hopping). Demultiplexing errors are ultimately caused by sequencing errors when the barcode sequence reported by the sequencer for a read has so many mistakes that the barcode sequence cannot be attibuted to the correct sample. In contrast, barcode crosstalk reflects an upstream failure in which an incorrect barcode becomes physically associated with a nucleic-acid fragment during library preparation or on-flowcell amplification. Barcode crosstalk masquerades as cross-contamination, which is particularly problematic for experiments that are sensitive to cross-contamination, such as those including low-biomass samples.

Substantial progress has been made in reducing demultiplexing error through improved barcode designs, basecalling and demultiplexing algorithms. For Oxford Nanopore Technologies (ONT) long-read sequencing, signal-level demultiplexers such as Deepbinner have improved assignment performance and reduced false assignments^1,2^. However this progress does nothing to mitigate barcode crosstalk, because barcode crosstalk results in the wrong label attached a nuleic-acid molecule.

Mitigating barcode crosstalk is essential for accuracy and reproducibility, because even low-level events can generate false positives, inflate diversity estimates, and distort quantitative analyses^3,4^. On Illumina patterned flow-cell instruments, for example, ExAmp-associated barcode crosstalk has been reported at rates ranging 0.1–6% depending on library type and indexing strategy^5,6^. Use of unique dual indexes (UDIs) is now standard practice to mitigate Illumina ExAmp-associated barcode crosstalk. However UDIs do nothing to reduce crosstalk, they are a mitigation strategy that allows identification and bioinformatic filtering, of crosstalked reads.

Here we identify substantial barcode crosstalk in multiplexed ONT sequence data generated with the Native Barcoding Kit. We also develop a framework to quantify this barcode crosstalk, and introduce post-ligation pooling (PLP), a simple workflow modification that substantially reduces crosstalk. We show that PLP reduces barcode misassignment by 1–2 orders of magnitude. Together, these results highlight the importance of considering the possibility of barcode crosstalk when analyzing ONT sequence data and provide an immediately adoptable solution.

## Results

### Real-sample metagenomics validates protocol improvements

We generated ONT shotgun metagenomic data from built-environment microbiome samples and observed what initially appeared to be cross-contamination. Negative controls (a molecular biology–grade water blank and a no-input empty blank) contained sequences consistent with both the p-trap water sample and the positive controls (*DCS lambda* and ΦX174), and the positive controls also contained sequences consistent with the environmental sample (Figure 1A). Because cross-contamination and background DNA are well-recognized challenges in microbiome studies, we evaluated common sources such as “kitome” and “splashome” contamination^3^ and found no procedural explanation consistent with the reciprocal signals observed across blanks, samples, and controls. We therefore investigated barcode crosstalk as a possible cause of the apparent cross-contamination.

**Figure 1.**
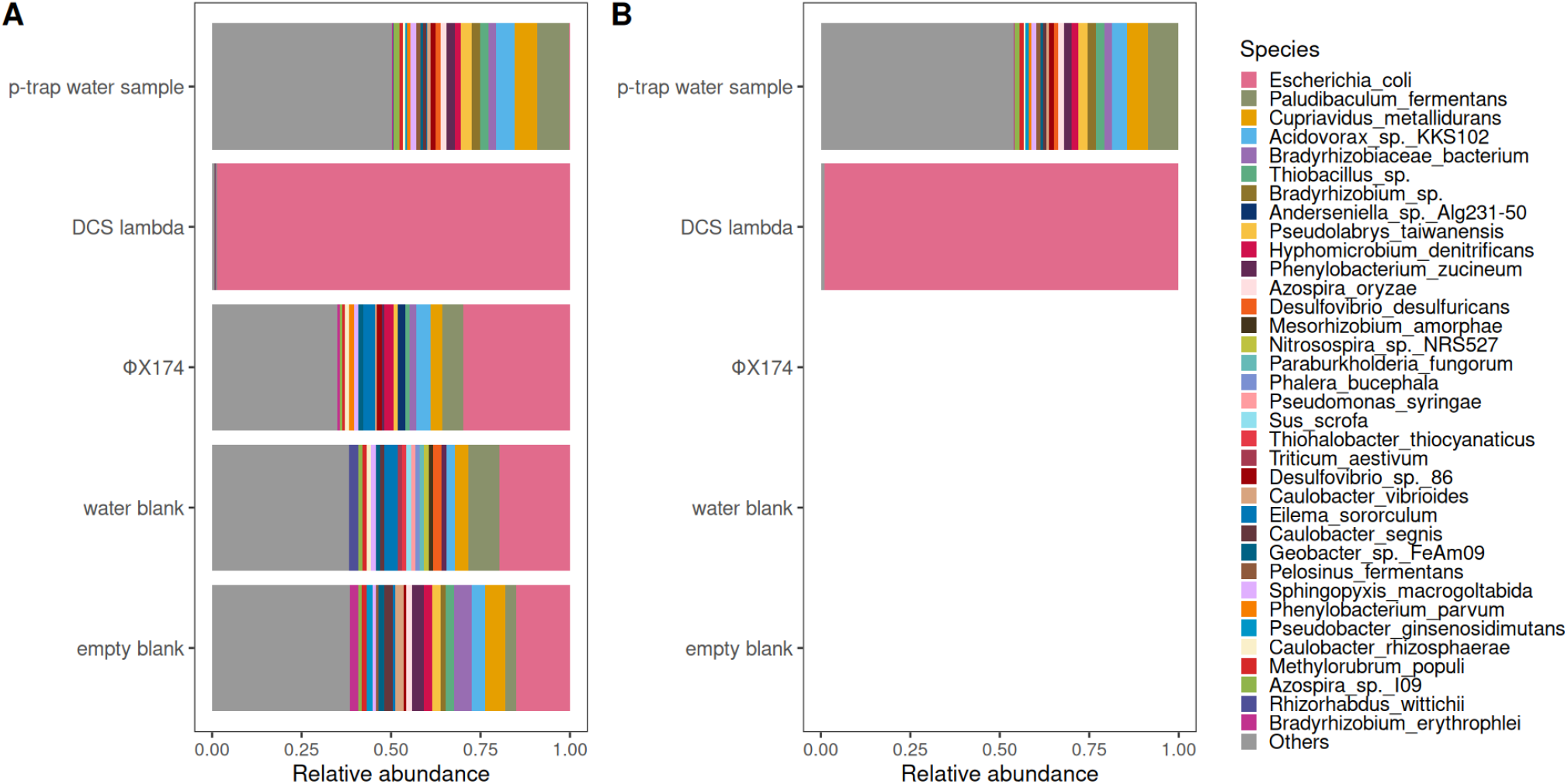
Taxonomic classification of libraries prepared with (A) standard protocol, and (B) modified PLP protocol. Note that reads from the spike-in phage controls are classified at the host level (*E. coli*) by this classifier/database.

We hypothesized that crosstalk was occurring during the adapter ligation step. In the standard ONT ligation workflow, samples are pooled immediately after index barcoding and prior to adapter ligation. We suspected that this pooled environment created an opportunity for excess index barcode to be ligated to DNA from any of the samples. To test this hypothesis, we developed PLP, a modified workflow in which libraries are pooled only after final adapter ligation. We found that PLP markedly reduced the apparent cross-contamination in the built-environment dataset (Figure 1B, Tables S1 and S2). Although there are still reads in the negative controls prepared by PLP, the numbers are 1-2 orders of magnitude lower than with the standard protocol (77 vs 1043 and 7 vs 1386 total reads;Table S1), and moreover none of these reads matches the MEGAN ^7^ taxonomy mapping (Figure 1B, Table S1), suggesting that these residual reads are non-microbial. A further analysis focusing on the positive controls found that PLP nearly eliminated (one *DCS lambda* read appears in the water blank) the appearance of positive control reads in other samples (Table S2). Note that ΦX174 is almost like a negative control here: very few reads are generated from ΦX174 genomic DNA because it is a double-stranded circular genome, so there are no ends to serve a substrate for ligation of barcode or adapter. The small number of ΦX174 sequences we observed were likely generated from ΦX174 DNA molecules with double-strand breaks.

To quantify the origin of the residual signal observed in negative controls under the standard workflow, we performed microbial source tracking using FEAST^8^. FEAST estimated that most taxonomically classified reads in standard-workflow blanks were attribuTable to the defined sources (FigureS1 and Table S3). In particular, FEAST attributed approximately 90% of classified reads in the water and empty blanks to the defined sources, consistent with barcode crosstalk being the dominant contributor to the apparent cross-contamination observed in negative controls. Together, the community profiles, absolute blank read counts, targeted mapping of positive controls, and source-tracking analysis indicate that PLP substantially reduces barcode crosstalk in a real-world context that is highly sensitive to contamination-like artifacts and improves interpretation of negative controls and positive controls.

### Quantification of Barcode Crosstalk

Our environmental data show that PLP substantially reduces barcode crosstalk in a real-world context sensitive to cross-contamination. However the complexity of the p-trap sample and the structure of the experiment limited our ability to quantify crosstalk. To more carefully quantify crosstalk-induced misassignment, we designed a minimal defined-genome experiment in which each barcode was associated with a simple, distinct DNA sample and the experiment was structured to avoid confounding by batch effects. DNA samples used were *DCS lambda* and three taxonomically distinct bacterial isolates (*Mycoplasma hominis, Phocaeicola vulgatus, Corynebacterium striatum*). Under this design, any read assigned to a barcode but classified or mapped to a different genome provides direct evidence of crosstalk-induced misassignment. We compared four library-preparation treatments: the standard ONT ligation protocol, PLP, ONT’s recently released Short Fragment Buffer (SFB) cleanup, and a combined SFB+PLP protocol. To avoid confounding protocol performance with sample or run-specific effects, we used a combinatorial, Latin-square–inspired design stratified by input amount. For each protocol comparison, the four DNA samples were partitioned into two sets, each assigned to one of the two protocols being compared; libraries from both protocols were then pooled at the adapter-cleanup step and sequenced on the same flow cell so that both protocols shared the same run conditions (Table S4). Across runs, the assignment of genomes to protocols was rotated so that no single genome systematically favored or disadvantaged any protocol, and differences in misassignment primarily reflect library preparation rather than run-to-run variability.

We quantified barcode crosstalk in two complementary ways. First, to compare the absolute burden of spurious reads across methods on a common scale, we report misassigned reads per million total reads, computed separately for DNA samples and water blanks within each library-preparation treatment (Figure 2). Second, to summarize overall assignment fidelity across samples, we computed recall, precision, F1, and the misassigned rate as a direct percentage using assigned mapped reads under each protocol (Table S5). These metrics are complementary: the per-million representation emphasizes the frequency of misassigned reads on an output-normalized scale, while the misassigned rate and F1 summarize overall assignment performance as percentages.

**Figure 2.**
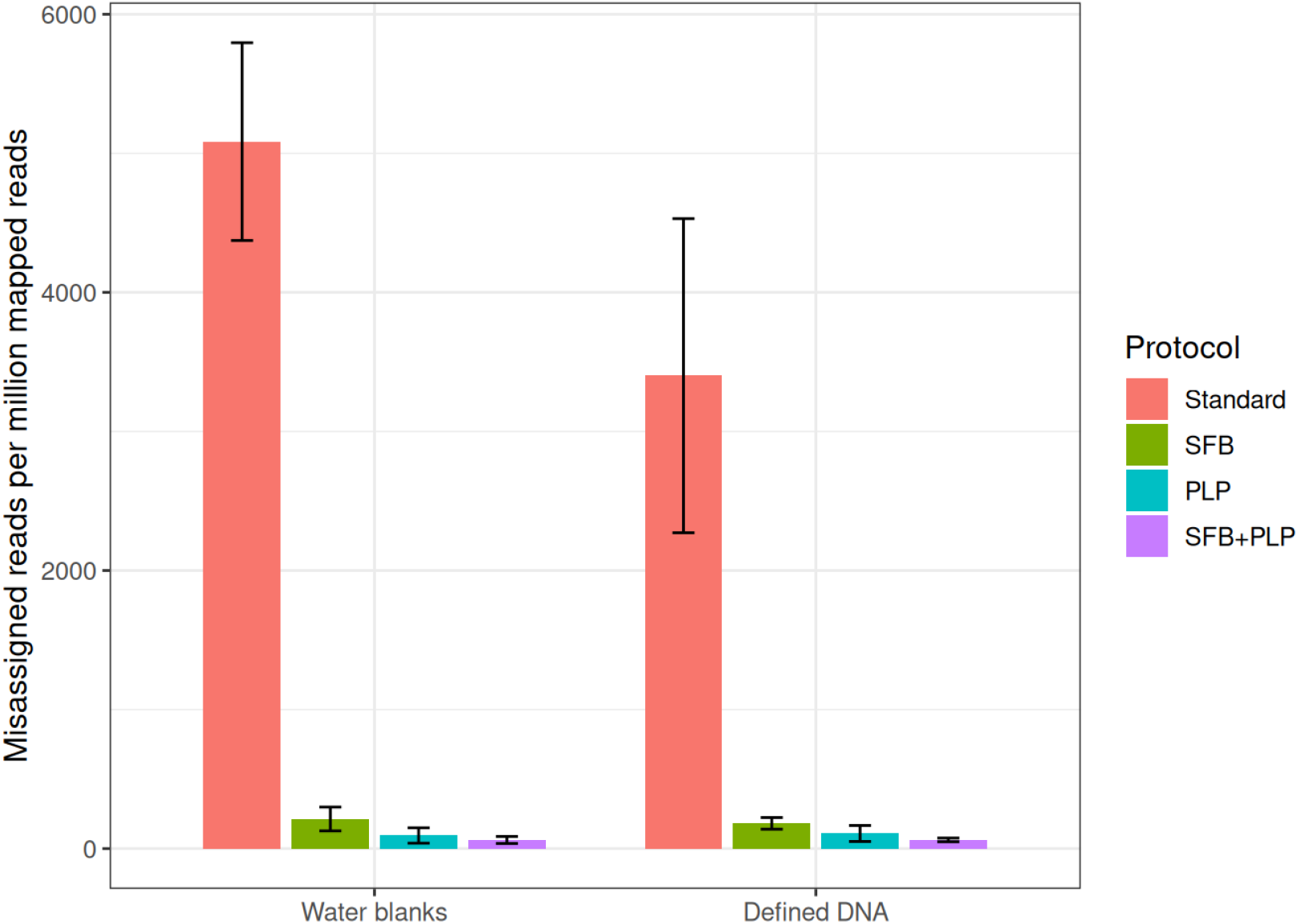
Misassigned reads per million mapped reads across different treatment.

Across experiments and sample types, the standard workflow produced substantially higher misassignment than the modified workflows (Figure 2; Table S5). Standard libraries showed misassignment at the level of thousands of reads per million in both defined DNA samples and water blanks, whereas SFB and PLP each reduced misassigned reads by approximately an order of magnitude, and the combined SFB+PLP workflow yielded the lowest misassignment across groups (Figure 2). When summarized across all barcoded samples using assigned, mapped reads (Table S5), the standard protocol exhibited the highest misassigned rate (0.882%), while SFB reduced misassignment to 0.067%, PLP reduced misassignment to 0.019%, and SFB+PLP achieved the lowest misassignment (0.015%). Together, these results show that deferring pooling until after adapter ligation (PLP), particularly in combination with SFB cleanup, substantially reduces crosstalk-induced misassignment across controlled single-genome libraries and negative controls.

## Discussion

Our results demonstrate high levels of barcode misassignment on the ONT platform when using the standard ONT native barcoding library preparation method. This level of barcode misassignment can materially affect results and interpretation, and should concern anyone using ONT native barcoding or analyzing data that was previously generated with this method. In our built-environment shotgun metagenomic experiment, the standard workflow produced blank and control profiles consistent with cross-contamination (Figure 1A), while PLP, our modified workflow, markedly suppressed these artifacts and reduced the absolute number of reads assigned to negative controls by 1–2 orders of magnitude (Figure 1B and Table S1). The substantial reduction of barcode misassignment when using our PLP method provides strong mechanistic evidence that the high levels of barcode misassignment with the standard method are a result of barcode crosstalk. It further suggests that barcode crosstalk is a result of unbarcoded DNA from multiple samples and free barcode experiencing a shared ligation environment. In the standard protocol, this shared ligation environment is possible because samples are pooled for adapter ligation (Figure 3, left).

**Figure 3.**
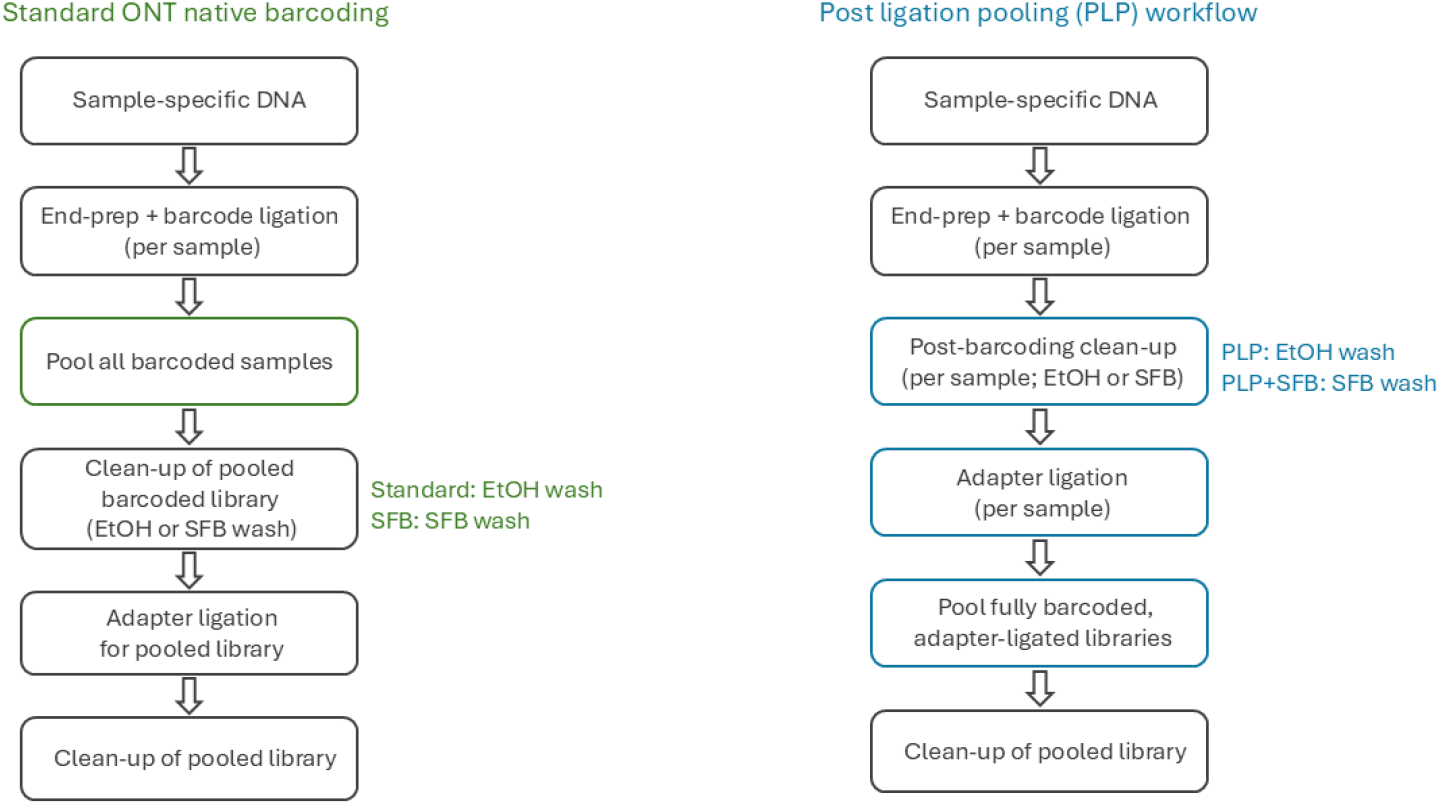
Comparison of pooling strategy in ONT native barcoding workflows

Our results with both SFB and PLP suggest that the ethanol cleanup after pooling is not sufficient to remove free barcode, and that this free barcode becomes ligated to unbarcoded sample DNA during the adapter ligation step. The SFB results demonstrate that barcode crosstalk can be substantially reduced by more effectively removing free barcode from the shared ligation environment. The PLP results demonstrate that barcode crosstalk can be substantially reduced by avoiding a shared ligation environment (Figure 3, right), which can be done by delaying sample pooling until after adapter ligation. Controlled defined-DNA experiments further quantified this effect, showing that protocol modifications—particularly PLP and SFB+PLP—reduce misassignment to 0.01–0.02% (Table S5), while preserving high overall assignment performance.

While this study is the first to identify, characterize, and remedy barcode crosstalk on the ONT platform, barcode crosstalk is a well know problem on Illumina sequencers that use ExAmp Cluster Amplification. Illumina-based pooled tagged-amplicon workflows can exhibit tag-jumping artifacts >2%^3^, and even a small barcode crosstalk rate of ∼0.3–0.9% on Illumina patterned flow cells can yield >8% barcode crosstalked molecules in low-input single-cell libraries^9^. In our ONT dataset, the standard protocol’s misassignment rate is of similar magnitude, a level unacceptable for applications where low-level signals can drive conclusions.

UDIs are the standard mitigation for Illumina barcode crosstalk. Incorporating UDIs into library prepartion enables crosstalked reads to be bioinformatically detected and excluded, reducing misassignment below 0.01%^6^. However, UDIs are a mitigation strategy, and they do not reduce barcode crosstalk. UDIs allow crosstalked reads to be bioinformatically detected and removed from downstream analysis, so crosstalk mitigation by UDI reduces the data usable for analysis. In contrast to UDIs, the crosstalk solutions we evaluate here have several desirable characteristics for ONT sequencing compared to the UDI approach because they are simple, highly effective, no-cost modifications. Most importantly, these solutions prevent crosstalk instead of simply mitigating it, which avoids wasting sequencing capacity on unusable reads. Furthermore these solutions acheive reduction in misassignment similar to UDIs: our PLP+SFB method reduced ONT misassignment to ∼0.01% (Table S5).

The solutions we evaluate here are particularly important for low-input or low-biomass contexts, where trace contamination and cross-contamination-like artifacts can dominate the measured signal^10^ (Figure 1A). Barcode crosstalk is especially problematic in such designs because it can introduce contamination-like artifacts into negative controls and low-abundance samples, inflating apparent diversity and biasing quantitative comparisons. By reducing barcode crosstalk, these solutions increase confidence that low-abundance reads reflect true sample content rather than misassignment.

More broadly, reducing barcode misassignment expands the suitability of ONT multiplexing for applications where cross-sample leakage can obscure weak or imbalanced signals. ONT sequencing is increasingly used in new applications as varied as simultaneous long-read profiling of somatic mutations and T-cell receptor repertoires^11^ and IgA-coating indexing^12^. With the PLP protocol (with or without SFB wash), libraries prepared from only a few nanograms of DNA can be multiplexed with greater confidence in intrinsically rare signals such as those in single-cell sequencing or environmental DNA detection in low-biomass microbiome samples. Finally we strongly encourage users of the ONT Native Barcoding Kit to immediately begin using one of the solutions to crosstalk that we evaluate here. We further encourage researchers analyzing data generated with the ONT Native Barcoding Kit to consider the possibility that crosstalk may be present in the data and evaluate the extent to which it could adversely affect their results. This is particularly urgent for research that is sensitive to cross-contamination or apparent cross-contamination.

## Methods

### Sequencing and Data Preprocessing

The Native Barcoding Kit (ONT, SQK-NBD114.96) was used for all DNA library preparation, with protocol modifications discussed below. A step by step online PLP protocol is available on https://www.protocols.io/private/509EBE88DD1411F0B60B0A58A9FEAC02for review.

POD5 files were basecalled and demultiplexed with Dorado v1.3.1^13^, after completion of sequencing, using a custom pipeline (https://github.com/QingDAI0225/dorado_basecalling_with_qscore). In addition to basecalling, the pipeline was used to extract per-read quality metrics and to perform read alignment using Dorado’s built-in Minimap2.

### Reference Genome Sequences and Databases

The *DCS lambda* sequence was downloaded from ONT https://a.storyblok.com/f/196663/x/f69b1ef376/dcs_reference.txt. The ΦX174 sequence was downloaded from NCBI (NC_001422.1), Genome sequences for ATCC 8482, ATCC 23114, and ATCC 6940 were downloaded from The ATCC genome portal^14^ (https://github.com/ATCC-Bioinformatics/genome_portal_api).

Taxonomy profiling used the BLAST nt database, downloaded from NCBI (last modified 2024-02-08;https://ftp.ncbi.nlm.nih.gov/blast/db/FASTA/nt.gz) and the MEGAN mapping file, downloaded from the MEGAN6 Download Page (megan-nucl-Feb2022;https://software-ab.cs.uni-tuebingen.de/download/megan6/megan-nucl-Feb2022.db.zip, as recommended in the instructions for the Taxonomic-Profiling-Minimap-Megan pipeline.

### Metagenomic Experiment and Analysis

Genomic DNA was extracted from a p-trap water sample using the QIAGEN DNeasy PowerSoil Pro Kit. The positive controls used were *DCS lambda* (Oxford Nanopore Technologies, DCS) and ΦX174 DNA (New England Biolabs, ΦX174 RF I DNA). Negative controls were a water blank (molecular biology–grade water substituted for DNA sample) and an “empty blank” (nothing was substituted for sample DNA during the library preparation). For each of these samples (p-trap, *DCS lambda*, ΦX174, water blank, and empty blank), we prepared two technical libraries from the same DNA: one using the PLP protocol and one using the standard protocol. The PLP and standard libraries were then sequenced on two separate Flongle flow cells, with 10 ng input DNA per sample (except negative controls) in all cases.

#### Taxonomy profiling

The Taxonomic-Profiling-Minimap-Megan pipeline ^15^, was used for taxonomy profiling, with minor adaptations for running on SLURM cluster.

#### Crosstalk Quantification

Crosstalk of positive control reads into the negative controls and p-trap sample was quantified by Minimap2.28^16^ using a custom reference constructed from the *DCS lambda* amplicon andΦX174 genome sequences.

#### Source tracking analysis

FEAST ^8^ was used to quantify how each sample contributed to the apparent cross-contamination due to barcode crosstalk. FEAST was run with EM_iterations = 1000 and with the setting in which multiple sinks are evaluated against the same donor set (different_sources_flag = 0). Species-level count Tables generated with the Minimap-MEGAN pipeline were the input to FEAST. For the purposes of source tracking, candidate “sinks” were the count Tables generated with the standard library preparation protocol from all samples (including blanks). Candidate “sources” were modified versions of the count Tables generated with the PLP protocol for the p-trap and *DCS lambda* samples. The source count Tables were modified as follows to account for the fact that the *E. coli* reference genome includes *lambda* prophage sequence, which results in the Minimap-MEGAN pipeline assigning all *DCS lambda* reads to *E. coli*: (1) In the PLP p-trap count Table all *E. Coli* counts were removed (i.e. *E. coli* counts were set to zero, counts for all other taxa were unmodified); (2) In the PLP *DCS lambda* count Table all counts other than *E. coli* were removed (i.e. *E. coli* was unmodified, counts for all other taxa were set to zero). We confirmed that the PLP p-trap water sample does not contain any *DCS lambda* reads by mapping PLP p-trap reads to the *DCS lambda* reference genome (Table S2).

### Quantification of Barcode Crosstalk

#### Experimental Design

The samples used for barcode crosstalk quantification were the bacterial isolates ATCC 8482 *Phocaeicola vulgatus*, ATCC 23114 *Mycoplasma hominis* and ATCC 6940 *Corynebacterium striatum*, obtained from ATCC as purified DNA and the positive control DNA (*DCS lambda* amplicon DNA provided in the ONT kit) were used.

Each treatment condition included two DNA samples and one blank (molecular biology–grade water) control and were run on both Flongle and MinION flowcells. To minimize batch effects resulting from variation in flow cell performance (e.g., available pores, active channel counts, or runtime stability), we implemented a combinatorial, Latin-square–inspired design stratified by input amount (Table S4). For each protocol comparison (standard vs PLP, SFB vs PLP, and SFB vs PLP+SFB), the four genomes were partitioned into two complementary two-sample sets, with sample-to-protocol assignments rotated across Flongle and MinION runs. Within each run, samples in one set were prepared using one protocol and samples in the other set were prepared using the other protocol. Libraries were pooled in the adapter-cleanup step and sequenced on the same flow cell.

#### Library Preparation and Sequencing

The standard preparation follows ONT’s Native Barcoding Kit protocol prior to Oct. 2025 (barcode, pool, ethanol wash, adapter ligation). The SFB preparation follows the ONT’s protocol released in Oct. 2025 (the standard protocol, but the ethanol wash is replaced with an SFB wash). PLP is the same as the Native Barcoding Kit protocol prior to Oct. 2025, but with pooling delayed until after adapter ligation. PLP+SFB follows the ONT’s protocol released in Oct. 2025, but again delays pooling until after adapter ligation. (Figure 3) Detailed step-by-step procedures are provided as Supplementary Information.

For each comparison, paired experiments were performed and sequenced on same flow cells: standard vs PLP, SFB vs PLP, and SFB vs SFB + PLP. In total, three MinION and three Flongle flow cells were used. Each Flongle library was prepared from 3 ng of input DNA per sample (12 ng in total), while each MinION library used 20 ng of input DNA per sample (80 ng in total).

#### Data Analysis

Dorado built-in alignment function was used to map raw reads to a “hybrid” reference, which was generated by concatenating the ATCC 8482, ATCC 23114, ATCC 6940 genomes and the *DCS lambda* amplicon sequences. No additional read-level quality or length filtering was applied. Only demultiplexed mapped reads were considered in calculating metrics (i.e. unclassified and unmapped reads were excluded). R was used for data analysis and Figure generation.

#### Barcode assignment performance metrics

We evaluated barcode-assignment performance using a confusion matrix constructed for each sequencing batch and library-preparation method. For each read, we recorded its barcode (sample) and its observed species assignment. Species assignments were restricted to the set of expected target species. Reads that did not map to any target genome, or that failed the mapQ threshold, were grouped into an “Unmapped” category for reporting.

For each batch–method–barcode combination, we tabulated the number of reads assigned to each target species (and to “Unmapped”). For DNA sample barcodes, the expected species for each barcode was known from the sample sheet. The count of reads assigned to the expected species for that barcode was treated as the “correctly assigned” count (true positives).

All barcode-level performance calculations were based on reads assigned to target species only (i.e., excluding the “Unmapped” reads).

#### Average precision/recall/F1

We computed barcode-level precision, recall, and F1 for each individual DNA sample barcode instance, and then summarized each method by averaging across these barcode instances. For a given DNA sample barcode instance, precision was calculated as the fraction of target-mapped reads from that barcode that were assigned to its expected species (correctly assigned reads divided by all target-mapped reads for that barcode). For a given DNA sample barcode instance, recall was calculated as the fraction of all reads assigned to the barcode’s expected species within the same batch that were attribuTable to that barcode. Importantly, the denominator for recall was computed within batch across all methods and including blank barcodes. This yields a conservative measure of species recovery when reads assigned to a species can be distributed across protocols within the same batch. For each barcode instance, we computed the F1 score as the harmonic mean of that instance’s precision and recall. For each library-preparation method, we report the average precision, average recall, and average F1 as the mean across all DNA sample barcode instances observed under that method across batches.

#### Misassigned rate by methods

In addition to barcode-averaged scores, we quantified the overall frequency of misassigned reads based on target-mapped reads: 1) For DNA sample barcodes, reads assigned to a non-expected target species were counted as misassigned; 2) For barcodes from blank samples, any read assigned to a target species was counted as misassigned (since blanks have no expected target species). Method-level misassigned rates were obtained by aggregating across batches using read-weighting (i.e., batches contributing more target-mapped reads contribute proportionally more to the final rate).

#### Misassigned reads per million and component plots

To summarize the absolute burden of barcode misassignment on a per-read basis, we converted misassignment counts to misassigned reads per million mapped reads using a shared denominator within each batch–protocol combination. All rates were based on reads mapped to the predefined target-species set (i.e., excluding the “Unmapped” category). For each batch and protocol combination, we defined 1) denominator is the combined total number of reads from DNA sample barcodes and blank barcodes that mapped to any target species; 2) numerator for DNA sample is reads from DNA sample barcodes mapped to a non-expected target species, and numerator for blank is reads from blank barcodes mapped to any target species. We then reported

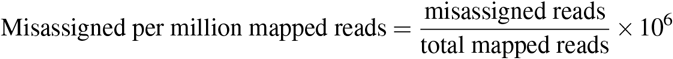

This calculation produced one value for each DNA sample within each batch–protocol combination, all normalized by the same shared denominator for that batch–protocol combination. Batch-resolved plots (Figure S2) were generated directly from these batch × method values, to visualize run-to-run variability.

Using the same denominator (total mapped reads within the batch×method), we split misassignment into two components: 1) Defined DNA: misassigned reads originating from DNA sample barcodes that mapped to non-expected target species. 2) Water blank: target-mapped reads observed in blank barcodes. Each component was expressed as per million mapped reads by dividing the component count by the same total mapped reads and multiplying by 10^6^. For protocol-level summaries (bar plot with error bars), we computed the mean across DNA sample barcode observations for each method, with error bars representing variability.

## Supporting information

Standard ligation sequencing protocol

PLP protocol

## Code Availability

All analyses are fully reproducible using the code available in the accompanying GitHub repository (https://doi.org/10.5281/zenodo.17981224), the publicly available Apptainer images indicated in the code, and the publicly available data available below.

## Data Availability

The sequence data have been deposited with links to BioProject accession number PRJNA1377585 in the NCBI BioProject database (https://www.ncbi.nlm.nih.gov/bioproject/).

## Acknowledgements

This work was supported by the Engineering Research Centers Program of the National Science Foundation under NSF Cooperative Agreement No. EEC-2133504. Any opinions, findings and conclusions or recommendations expressed in this material are those of the author(s) and do not necessarily reflect those of the National Science Foundation.

## Author contributions statement

QD: conceptualization, methodology, software, validation, formal analysis, investigation, data curation, writing - original draft, writing - review & editing, visualization;

CKG: resources, writing - review & editing, supervision, project administration, funding acquisition;

JAG: software, visualization, resources, writing - review & editing, supervision, project administration, funding acquisition.

