## Supplementary material for "Identification, quantification, and elimination of barcode crosstalk in multiplexed Oxford Nanopore sequencing": Supplimentary Tables and Figures

### Supplementary Information

**Table S1.** Species-level read statistics for each sample and protocol in case study.

| Sample | Protocol | Total reads | Species-level reads | Species-level fraction |
|---|---|---|---|---|
| p-trap water sample | PLP | 39456 | 5301 | 0.134 |
| p-trap water sample | standard | 196865 | 23702 | 0.120 |
| *DCS lambda* | PLP | 2878 | 1484 | 0.516 |
| *DCS lambda* | standard | 8314 | 3994 | 0.480 |
| ΦX174 | PLP | 143 | 0 | 0.000 |
| ΦX174 | standard | 2333 | 154 | 0.066 |
| water blank | PLP | 77 | 0 | 0.000 |
| water blank | standard | 1043 | 81 | 0.078 |
| empty blank | PLP | 7 | 0 | 0.000 |
| empty blank | standard | 1386 | 127 | 0.092 |

**Table S2.** Number of reads mapped to ΦX174 genome and *DCS lambda* reference sequence by Minimap2.

| Sample | Protocol | *DCS lambda* Reads | ΦX174 Reads | Total reads |
|---|---|---|---|---|
| p-trap water sample | PLP | 0 | 0 | 39456 |
| p-trap water sample | standard | 61 | 5 | 196869 |
| *DCS lambda* | PLP | 2814 | 0 | 2978 |
| *DCS lambda* | standard | 7320 | 0 | 8568 |
| ΦX174 | PLP | 0 | 107 | 193 |
| ΦX174 | standard | 49 | 641 | 2657 |
| water blank | PLP | 1 | 0 | 77 |
| water blank | standard | 32 | 1 | 1044 |
| empty blank | PLP | 0 | 0 | 7 |
| empty blank | standard | 38 | 4 | 1389 |

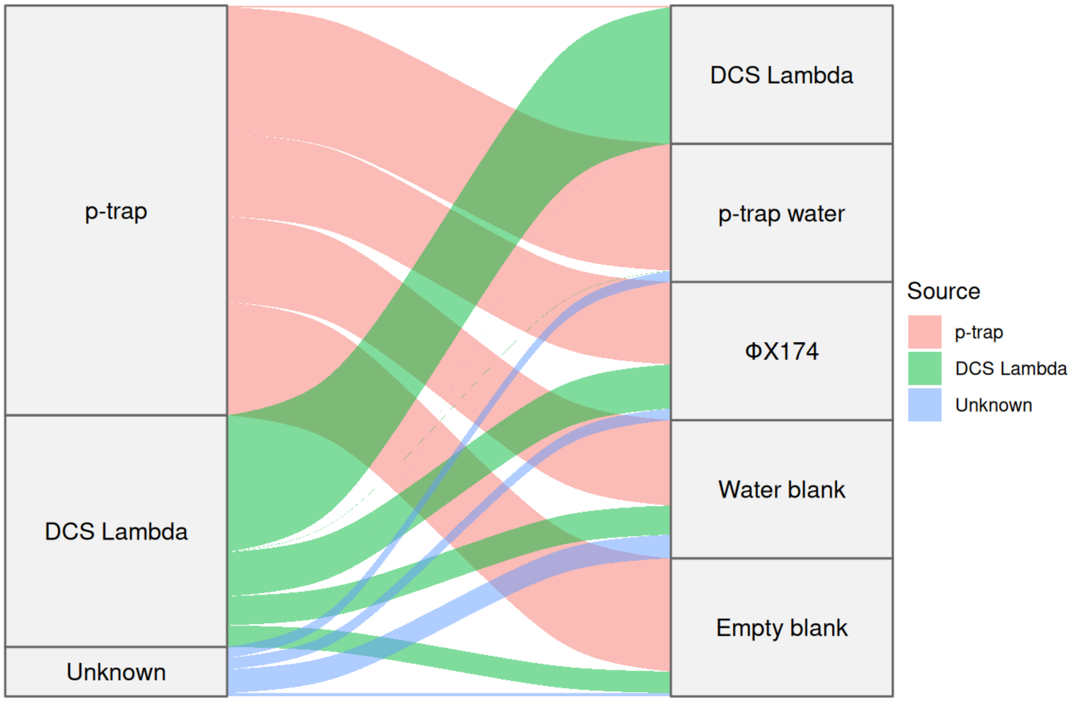


**Figure S1.** FEAST source tracking attributes apparent contamination of blanks signal under the standard workflow to p-trap and *DCS lambda*.

**Table S3. FEAST source attribution.** Numerical results from FEAST source attribution used to generate Figure S1.

| Sample in standard workflow | p-trap | *DCS lambda* | Unknown |
|---|---|---|---|
| p-trap | 0.9473 | 0.0021 | 0.0507 |
| *DCS lambda* | 0.0095 | 0.9879 | 0.0027 |
| ΦX174 | 0.5333 | 0.3067 | 0.1600 |
| water blank | 0.6538 | 0.2051 | 0.1410 |
| empty blank | 0.7705 | 0.1557 | 0.0738 |

**Table S4.** Summary of pairwise comparisons for different input amounts and protocols. The pairwised protocol samples are pooled together in adapter cleaning step and sequenced on the same flow cell. The single-letter codes indicate the species of input DNA: M = Mycoplasma hominis, P = Phocaeicola vulgatus, C = Corynebacterium striatum, and L = *DCS lambda*.

| | standard vs PLP | | SFB vs PLP | | SFB vs PLP+SFB | |
|---|---|---|---|---|---|---|
| | standard | PLP | SFB | PLP | SFB | PLP+SFB |
| 3ng Flongle | ML | PC | MP | CL | LC | PM |
| 20ng MinION | MP | CL | CP | LM | ML | PC |

**Table S5.** Performance of different library prep treatments, only assigned mapped reads included.

| Method | Mean Recall (%) | Mean Precision (%) | Mean F1 (%) | Misassigned rate (%) | number of samples |
|---|---|---|---|---|---|
| standard | 98.705 | 99.229 | 98.966 | 0.882 | 4 |
| SFB | 99.892 | 99.963 | 99.928 | 0.067 | 8 |
| PLP | 99.983 | 99.949 | 99.966 | 0.019 | 8 |
| SFB+PLP | 99.994 | 99.985 | 99.990 | 0.015 | 4 |

**Table S6.** Confusion matrix for Flongle flow cell with 3 ng input DNA for each DNA sample comparing Standard and PLP protocols.

| Sample | Protocol | *DCS lambda* | Mycoplasma hominis | Corynebacterium striatum | Phocaeicola vulgatus | Unmapped |
|---|---|---|---|---|---|---|
| L (3 ng) | standard | 1950 | 29 | 0 | 0 | 1959 |
| M (3 ng) | standard | 34 | 3906 | 0 | 0 | 391 |
| Blank | standard | 14 | 12 | 0 | 0 | 72 |
| C (3 ng) | PLP | 1 | 0 | 624 | 1 | 56 |
| P (3 ng) | PLP | 0 | 0 | 0 | 3311 | 69 |
| Blank | PLP | 0 | 1 | 0 | 0 | 30 |

**Table S7.** Confusion matrix for Flongle flow cell with 3 ng input DNA for each DNA samples comparing SFB and PLP protocols.

| Sample | Protocol | Mycoplasma hominis | Phocaeicola vulgatus | *DCS lambda* | Corynebacterium striatum | Unmapped |
|---|---|---|---|---|---|---|
| M (3ng) | SFB | 13827 | 6 | 0 | 0 | 3236 |
| P (3ng) | SFB | 1 | 13279 | 0 | 0 | 264 |
| Blank | SFB | 0 | 0 | 0 | 0 | 70 |
| L (3ng) | PLP | 0 | 1 | 9305 | 0 | 1134 |
| C (3ng) | PLP | 0 | 1 | 0 | 9044 | 1838 |
| Blank | PLP | 0 | 0 | 0 | 0 | 284 |

**Table S8.** Confusion matrix for Flongle flow cell with 3 ng input DNA for each DNA sample comparing SFB and SFB + PLP protocols.

| Sample | Protocol | *DCS lambda* | Corynebacterium striatum | Phocaeicola vulgatus | Mycoplasma hominis | Unmapped |
|---|---|---|---|---|---|---|
| L (3ng) | SFB | 46859 | 21 | 0 | 0 | 1302 |
| C (3ng) | SFB | 18 | 25336 | 1 | 0 | 638 |
| Blank | SFB | 15 | 8 | 0 | 0 | 399 |
| P (3ng) | SFB + PLP | 4 | 0 | 46397 | 0 | 703 |
| M (3ng) | SFB + PLP | 6 | 0 | 0 | 22392 | 1086 |
| Blank | SFB + PLP | 6 | 0 | 0 | 0 | 105 |

**Table S9.** Confusion matrix for MinION flow cell with 20 ng input DNA for each DNA sample comparing SFB and SFB + PLP protocols.

| Sample | Protocol | Mycoplasma hominis | *DCS lambda* | Phocaeicola vulgatus | Corynebacterium striatum | Unmapped |
|---|---|---|---|---|---|---|
| M (20 ng) | SFB | 302875 | 117 | 1 | 3 | 13850 |
| L (20 ng) | SFB | 673 | 1802396 | 37 | 6 | 18849 |
| Blank | SFB | 456 | 342 | 1 | 3 | 1011 |
| P (20 ng) | SFB + PLP | 3 | 41 | 484962 | 3 | 5083 |
| C (20 ng) | SFB + PLP | 0 | 13 | 3 | 112879 | 1704 |
| Blank | SFB + PLP | 2 | 20 | 0 | 0 | 593 |

**Table S10.** Confusion matrix for MinION flow cell with 20 ng input DNA for each DNA sample comparing SFB and PLP protocols.

| Sample | Protocol | Corynebacterium striatum | Phocaeicola vulgatus | *DCS lambda* | Mycoplasma hominis | Unmapped |
|---|---|---|---|---|---|---|
| C (20 ng) | SFB | 621345 | 119 | 39 | 6 | 7308 |
| P (20 ng) | SFB | 29 | 231258 | 16 | 0 | 2545 |
| Blank | SFB | 14 | 56 | 60 | 1 | 4530 |
| L (20 ng) | PLP | 22 | 35 | 877685 | 88 | 35494 |
| M (20 ng) | PLP | 30 | 62 | 183 | 2428826 | 110750 |
| Blank | PLP | 25 | 84 | 102 | 19 | 12041 |

**Table S11.** Confusion matrix for MinION flow cell with 20 ng input DNA for each DNA sample comparing Standard and PLP protocols.

| Sample | Protocol | Phocaeicola vulgatus | Mycoplasma hominis | *DCS lambda* | Corynebacterium striatum | Unmapped |
|---|---|---|---|---|---|---|
| P (20 ng) | standard | 592049 | 3508 | 103 | 19 | 40180 |
| M (20 ng) | standard | 1673 | 1182259 | 69 | 9 | 63471 |
| Blank | standard | 2555 | 7721 | 85 | 13 | 14564 |
| L (20 ng) | PLP | 24 | 60 | 1067627 | 9 | 18560 |
| C (20 ng) | PLP | 8 | 23 | 36 | 201889 | 5867 |
| Blank | PLP | 9 | 9 | 46 | 4 | 4625 |

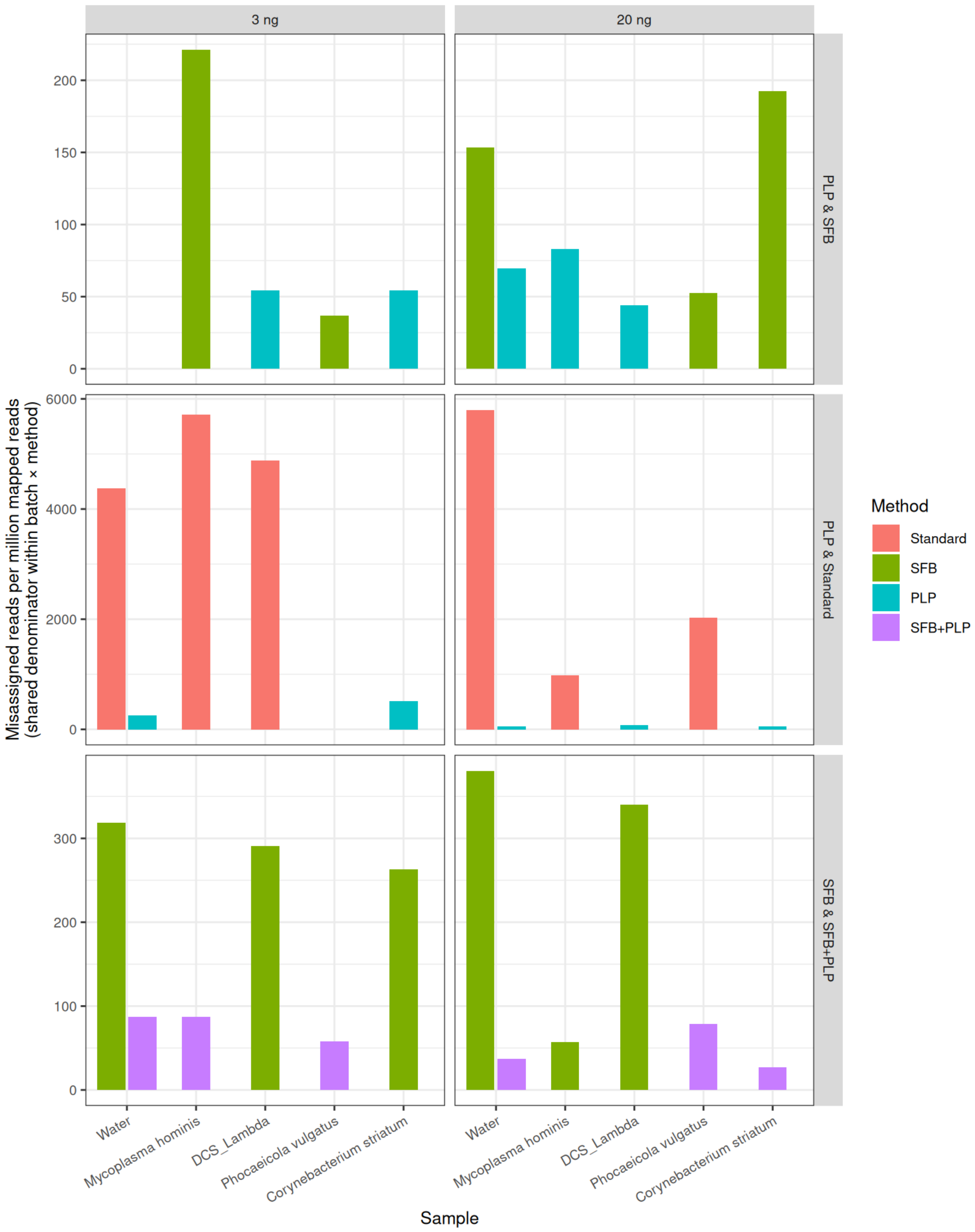


**Figure S2.** Misassigned read rates in each sequencing batch, stratified by input DNA amount and method. Each panel compares two methods' misassignment frequencies (per million reads) across the five sample types in a single batch. Left: 3 ng input libraries; Right: 20 ng input. Bottom: standard (red) vs. PLP (blue) – highlighting the large drop in crosstalk when adopting PLP at both low and high input. Middle: SFB (green) vs. SFB+PLP (purple) – showing the added benefit of PLP even with SFB cleanup (note the C. striatum bar in the 20 ng SFB run, which PLP mitigates). Top: SFB (green) vs. PLP (blue) – comparing the two strategies individually. In all cases, PLP-based preparations yield significantly fewer misassigned reads than the legacy standard approach, regardless of input quantity.
