## Supplementary material for "Identification, quantification, and elimination of barcode crosstalk in multiplexed Oxford Nanopore sequencing": Standard ligation sequencing protocol

### SQK-NBD114 High Yield Protocol

#### Overview

This protocol adapts an existing Oxford Nanopore technical document with a single goal: maximize data yield when starting material is below ONT's recommended input amounts. In prior workflows, mismatches between upstream and downstream steps could cause only a small fraction of the input material to progress into sequencing. Here, we streamline and harmonize step interfaces to minimize losses at transfers/cleanups and ensure that all prepared samples advance to the final sequencing stage. The procedure is explicitly engineered for sub-recommended inputs, i.e., any low-biomass or diluted samples, without changing the core chemistry, making it a drop-in, yield-focused improvement for challenging inputs.

Here's a reference-card style protocol tailored for MinION, 20 ng input per sample with SQK-NBD114.96, adapted from ONT NBE_9171_v114_revT_03Oct2025. I've highlighted our modifications so you can see exactly what changed.

#### Materials (as per ONT document unless noted)

- SQK-NBD114.96 kit (barcodes, ligation reagents, adapters)
- NEBNext® FFPE DNA Repair Mix (NEB, M6630), NEBNext® Ultra™ II End Repair/dA-Tailing Module (NEB, E7546)
- Nuclease-free water (e.g. ThermoFisher, AM9937)
- SFB Expansion (EXP-SFB001) or molecular biology grade ethanol (see note below).
- 1.5 ml Eppendorf DNA LoBind tubes, Eppendorf twin.tec® PCR plate 96 LoBind, semi skirted (Merck, EP0030129504) with heat seals
- P1000, P200, P20 and P10 pipette and tips
- Vortex mixer
- Hula mixer or rotator mixer
- Thermal cycler
- Ice bucket with ice
- Magnetic rack
- Qubit (optional for spot checks)

**Note on SFB vs ethanol (EtOH)**
Before ONT's revT protocol, the default post barcoding cleanup used 80% ethanol. In revT version, ONT changed to SFB washes after barcoding pooling and indicates SFB Expansion has to be purchased for this step. We evaluated both versions (pre-revT ethanol workflow, i.e. Standard protocol, and revT with SFB, i.e. SFB protocol). Yield were comparable when EtOH washes were done carefully.

#### 1. DNA repaire and end-prep (~20 minutes)

Prepare the NEBNext FFPE DNA Repair Mix and NEBNext Ultra II End Repair / dA tailing Module reagents in accordance with manufacturer's or Nanopore's original instructions, and place on ice.

In a clean 96-well plate, aliquot each sample and negative control. Make up each sample to **6** $\mu l$ using nuclease-free water. Mix gently by pipetting and spin down.

Combine the following components per well:

| Reagent | Volume |
|---|---|
| DNA sample | 6 $\mu l$ |
| NEBNext FFPE DNA Repair Buffer | 0.4375 $\mu l$ |
| Ultra II End-prep Reaction Buffer | 0.4375 $\mu l$ |
| Ultra II End-prep Enzyme Mix | 0.375 $\mu l$ |
| NEBNext FFPE DNA Repair Mix | 0.25 $\mu l$ |
| Total | 7.5 $\mu l$ |

**Note on volume modification**
To mitigate dead-volume losses, we use half (0.5×) of ONT's recommended volumes for all reagents while maintaining the same stoichiometries specified by ONT.

Note: We recommend mixing NEBNext FFPE DNA Repair Buffer, Ultra II End-prep Reaction Buffer, Ultra II End-prep Enzyme Mix and NEBNext FFPE DNA Repair Mix as a master mix, thoroughly mixed by pipetting, and add 1.5 $\mu l$ of the master mix for each well to simplify the loading of this step.

Between each addition, pipette mix 10-20 times.

Ensure the components are thoroughly mixed by pipetting and spin down briefly.

Using a thermal cycler, incubate at 20°C for 5 minutes and 65°C for 5 minutes.

#### 2. Native barcoding ligation (~60 minutes)

Prepare the NEB Blunt/TA Ligase Master Mix according to the manufacturers instructions, and place on ice.

Thaw the AMPure XP Beads (AXP) at room temperature and mix by vortexing. Keep the beads at room temperature.

Thaw the EDTA at room temperature and mix by vortexing. Then spin down and place on ice.

If you plan to use SFB in the washing step, thaw SFB at room temperature and mix by vortexing. Place on ice.

Thaw the Native Barcodes (NB01-96) required for your number of samples at room temperature. Individually mix the barcodes by pipetting, spin down, and place them on ice.

Select a unique barcode for every sample to be run together on the same flow cell. Up to 96 samples can be barcoded and combined in one experiment. Only use one barcode per sample.

In a new 96-well plate, add the reagents in the following order per well mixing well by pipetting between each addition:

| Reagent | Volume |
|---|---|
| End-prepped DNA | 7.5 $\mu l$ |
| Native Barcode (NB01-96) | 2.5 $\mu l$ |
| Blunt/TA Ligase Master Mix | 10 $\mu l$ |
| Total | 20 $\mu l$ |

**Note on volume modification**
In this step, we doubled each reagent in the reaction system to make sure all 7.5 $\mu l$ of end-prepped DNA go to the barcoding and following step.

Thoroughly mix the reaction by gently pipetting and briefly spinning down.

Incubate for 20 minutes at room temperature.

Add 4 $\mu l$ EDTA (blue cap) or 2 $\mu l$ EDTA (clear cap) to each well and mix thoroughly by pipetting and spin down briefly.

Pool the barcoded samples in a 1.5 ml Eppendorf DNA LoBind tube.

| | Volume per sample | For 24 samples | For 48 samples |
|---|---|---|---|
| Total volume for preps using EDTA (blue cap) | 24 $\mu l$ | 576 $\mu l$ | 1152 $\mu l$ |

Resuspend the AMPure XP Beads (AXP) by vortexing.

Add 0.4X AMPure XP Beads (AXP) to the pooled reaction, and mix by pipetting.

| | Volume per sample | For 24 samples | For 48 samples |
|---|---|---|---|
| Volume of AXP for preps using EDTA (blue cap) | 9.6 $\mu l$ | 230.4 $\mu l$ | 460.8 $\mu l$ |

Incubate on a Hula mixer (rotator mixer) for 10 minutes at room temperature.

Spin down the sample and pellet on a magnet for 5 minutes. Keep the plate on the magnetic rack until the eluate is clear and colourless, and pipette off the supernatant.

Wash the beads with 700 µl of Short Fragment Buffer (SFB). Flick the beads to resuspend, spin down, then return the sample to the magnetic rack and allow the beads to pellet. Remove the buffer using a pipette and discard.

Repeat the previous step.

Spin down and place the tube back on the magnetic rack. Pipette off any residual buffer.

Remove the tube from the magnetic rack and resuspend the pellet in 32 $\mu l$ nuclease-free water by gently flicking.

**Note on volume modification**
Note: Here we modified 35 $\mu l$ into 32 $\mu l$ because the next step requires 30 $\mu l$ of pooled barcoded sample. For those who want to check DNA concentration after washing, we leave 1 $\mu l$ available for optional Qubit quantification after the washing step while maintaining sufficient volume for the next reaction. Therefore, 32 $\mu l$ is selected to maximize recovery/yield without over-diluting the library.

Incubate for 10 minutes at 37°C. Every 2 minutes, agitate the sample by gently flicking for 10 seconds to encourage DNA elution.

Pellet the beads on a magnetic rack until the eluate is clear and colourless.

Remove and retain 32 $\mu l$ of eluate into a clean 1.5 ml Eppendorf DNA LoBind tube.

#### 3. Adapter ligation and clean-up (~50 minutes)

All steps are the same as Nanopore's recommended.

#### 4. Priming and loading the MinION and GridION Flow Cell (~10 minutes)

All steps are the same as Nanopore's recommended.
