## Supplementary material for "Identification, quantification, and elimination of barcode crosstalk in multiplexed Oxford Nanopore sequencing": PLP protocol

### SQK-NBD114 Post Ligation Pooling (PLP) High Yield Protocol

#### Overview

This protocol is built on top of the high-yield version (see SQK-NBD114 high yield protocol) and specifically adjusts the pooling strategy to reduce barcode crosstalk. In the original workflow, samples were pooled immediately after barcode addition and EDTA quench, followed by shared cleanup and adapter ligation. In this modified version, each sample is cleaned individually after barcode + EDTA, then kept separate for per-sample adapter ligation. Only after all ligation steps are complete are the libraries pooled, followed by a final SFB cleanup step. This preserves the yield benefits of the high-input–agnostic protocol while physically separating ligation reactions, thereby reducing opportunities for barcode misassignment and cross-ligation between samples.

Here's a reference-card style protocol tailored for MinION, 20 ng input per sample with SQK-NBD114.96, adapted from high yield version of ONT NBE_9171_v114_revT_03Oct2025. I've highlighted our modifications vs original ONT protocol so you can see exactly what changed.

#### Materials (as per ONT document unless noted)

- SQK-NBD114.96 kit (barcodes, ligation reagents, adapters)
- NEBNext® FFPE DNA Repair Mix (NEB, M6630), NEBNext® Ultra™ II End Repair/dA-Tailing Module (NEB, E7546)
- Nuclease-free water (e.g. ThermoFisher, AM9937)
- SFB Expansion (EXP-SFB001) or molecular biology grade ethanol (see note below).
- 1.5 ml Eppendorf DNA LoBind tubes, Eppendorf twin.tec® PCR plate 96 LoBind, semi skirted (Merck, EP0030129504) with heat seals
- P1000, P200, P20 and P10 pipette and tips
- Vortex mixer
- Hula mixer or rotator mixer
- Thermal cycler
- Ice bucket with ice
- Magnetic rack
- Qubit (optional for spot checks)

**Note on SFB vs ethanol (EtOH)**
Before ONT's revT protocol, the default post barcoding cleanup used 80% ethanol. In revT version, ONT changed to SFB washes after barcoding pooling and indicates SFB Expansion has to be purchased for this step. We evaluated both versions (pre-revT EtOH workflow, i.e. PLP protocol) and revT with SFB, i.e. PLP + SFB protocol). Yield were comparable when EtOH washes were done carefully.

Add 0.4x AMPure XP Beads (AXP) to each reaction, and mix by pipetting.

**Note on separate washing**
We add AMPure XP Beads on each sample. For example, if you use blue cap EDTA, you sample volume is 24 $\mu l$, and you would add 9.6 $\mu l$ AXP Beads to each sample. Otherwise, if you use clear cap EDTA, you would add 8.8 $\mu l$ AXP Beads to each sample.

Incubate on a Hula mixer (rotator mixer) for 10 minutes at room temperature.

Spin down the sample and pellet on a magnet for 5 minutes. Keep the plate on the magnetic rack until the eluate is clear and colourless, and pipette off the supernatant.

Wash the beads with 115 $\mu l$ of Short Fragment Buffer (SFB). Flick the beads to resuspend, spin down, then return the sample to the magnetic rack and allow the beads to pellet. Remove the buffer using a pipette and discard.

**Note on SFB volume**
In this step, we use 115 $\mu l$ SFB/EtOH per sample for the wash step because the original protocol specifies a single 700 $\mu l$ SFB/EtOH wash on the pooled library. In our example, we process 6 samples, so 115 $\mu l \times 6 \approx 700$ $\mu l$, keeping the total SFB/EtOH usage comparable to the original method while performing washes separately. For other batch sizes, the SFB/EtOH volume per sample can be adjusted proportionally so that the sum across all samples approximates the 700 $\mu l$ used in the original protocol.

Repeat the previous step.

Spin down and place the tube back on the magnetic rack. Pipette off any residual buffer.

Remove the tube from the magnetic rack and resuspend the pellet in 17 $\mu l$ nuclease-free water by gently flicking.

**Note on volume modification**
Note: Here we modified 35 $\mu l$ into 17 $\mu l$ because the next step requires 15 $\mu l$ for each barcoded sample. For those who want to check DNA concentration after washing, we leave 1 $\mu l$ available for optional Qubit quantification after the washing step while maintaining sufficient volume for the next reaction. Therefore, 17 $\mu l$ is selected to maximize recovery/yield without over-diluting the library.

Incubate for 10 minutes at 37°C. Every 2 minutes, agitate the sample by gently flicking for 10 seconds to encourage DNA elution.

Pellet the beads on a magnetic rack until the eluate is clear and colourless.

Remove and retain 17 $\mu l$ of eluate into a clean 1.5 ml Eppendorf DNA LoBind tube.

#### 3. Adapter ligation and clean-up (~50 minutes)

Prepare the NEBNext Quick Ligation Reaction Module according to the manufacturer's or ONT's instructions, and place on ice.

Spin down the Native Adapter (NA) and Quick T4 DNA Ligase, pipette mix and place on ice.

Thaw the Elution Buffer (EB) at room temperature and mix by vortexing. Then spin down and place on ice.

Thaw either Long Fragment Buffer (LFB) or Short Fragment Buffer (SFB) at room temperature and mix by vortexing. Place on ice.

Prepare diluted NA based on number of samples in your library as follows:

| Reagent | Volume for 2 samples | for 12 samples | for 24 samples |
|---|---|---|---|
| Original NA | 5 $\mu l$ | 5 $\mu l$ | 5 $\mu l$ |
| Molecular grade water | 0 $\mu l$ | 25 $\mu l$ | 55 $\mu l$ |
| Total | 5 $\mu l$ | 30 $\mu l$ | 60 $\mu l$ |

In 1.5 ml Eppendorf LoBind tube for each sample, mix in the following order: Between each addition, pipette mix 10-20 times.

| Reagent | Volume |
|---|---|
| Barcoded sample | 15 $\mu l$ |
| Diluted NA | 2.5 $\mu l$ |
| NEBNext Quick Ligation Reaction Buffer (5X) | 5 $\mu l$ |
| Quick T4 DNA Ligase | 2.5 $\mu l$ |
| Total | 25 $\mu l$ |

**Note on adapter dilution and volume modification**
We halve the reaction volumes at this stage because pooling is intentionally deferred until after adapter ligation to mitigate barcode crosstalk. Without volume reduction, keeping all samples separate through ligation would lead to an excessively large total volume at the subsequent pooling step. To avoid this while maintaining performance, we scale all components to 0.5× of the ONT-recommended volumes, preserving the original reagent ratios so that reaction stoichiometry and product quality are not affected. For the adapter, the original protocol specifies 5 $\mu l$ per batch; in this modified workflow, we dilute and redistribute the adapter so that the effective adapter amount per sample matches the reduced reaction volume and remains within ONT's intended range. This prevents overloading the library with excess adapter, which could otherwise occupy nanopores as free adapter complexes and reduce the fraction of productive, library-derived reads during sequencing.

**Note**
We recommend mixing diluted NA and NEBNext Quick Ligation Reaction Buffer (5X) as a master mix, thoroughly mixed by pipetting, and add 7.5 $\mu l$ of the master mix for each tube to simplify the loading of this step.

Incubate the reaction for 20 minutes at room temperature.

Pool all samples together.

Add 0.4x AMP beads, and mixed by pipetting.

All downstream steps are the same as Nanopore's recommended.

#### 4. Priming and loading the MinION and GridION Flow Cell (~10 minutes)

All steps are the same as Nanopore's recommended.
